# Forage stoichiometry predicts the home range size of a small terrestrial herbivore

**DOI:** 10.1101/2020.08.13.248831

**Authors:** Matteo Rizzuto, Shawn J. Leroux, Eric Vander Wal, Isabella C. Richmond, Travis R. Heckford, Juliana Balluffi-Fry, Yolanda F. Wiersma

## Abstract

Consumers make space use decisions based on resource quality. Most studies that investigate the influence of resource quality on the spatial ecology of consumers use diverse proxies for quality including measures based on habitat classification, forage species diversity and abundance, and nutritional indicators, e.g., protein. Ecological stoichiometry measures resource quality in terms of elemental ratios, e.g., carbon (C):nitrogen (N) ratio, but rarely have these currencies been used to study consumer space use decisions. Yet, elemental ratios provide a uniquely quantitative way to assess resource quality. Consequently, ecological stoichiometry allows for investigation of how consumers respond to spatial heterogeneity in resource quality by changing their space use, e.g. their home range size, and how this may influence ecosystem dynamics and trophic interactions. Here, we test whether the home range size of a keystone boreal herbivore, the snowshoe hare (*Lepus americanus*), varies with differences in the C:N, C:phosphorus (P), and N:P ratios of two preferred forage species, lowland blueberry (*Vaccinium angustifolium*) and red maple (*Acer rubrum*). We consider forage resources with higher C content relative to N and P to be lower quality than resources with lower relative C content. We use a novel approach, combining elemental distribution models with herbivore home range size estimates to test our hypothesis that hare home range size will be smaller in areas with access to high, homogeneous resource quality compared to areas with access to low, heterogeneous resource quality during summer months. Our results support our prediction for lowland blueberry, but not for red maple. Herbivore home range size decreased with increasing blueberry foliage quality, but also with decreasing spatial heterogeneity in blueberry foliage quality, i.e. N or P content. Herbivores in the boreal forest face strong nutritional constraints due to the paucity of N and P. Access to areas of high, homogeneous resource quality is paramount to meeting their dietary requirements with low effort. In turn, this may influence community (e.g., trophic interactions) and ecosystem (e.g., nutrient cycling) processes. Paradoxically, our study shows that taking a reductionist approach of viewing resources through a biochemical lens can lead to holistic insights of consumer spatial ecology.

## Introduction

Environmental and organismal variability within ecosystems are tightly interconnected. Geochemical, atmospheric, and biological factors drive differences in the elemental composition of primary producers across landscapes (Ågren and Weih, 2012; Borer et al., 2015; He et al., 2015). For example, elemental ratios in marine phytoplankton can vary widely across latitudinal gradients of environmental variables (e.g., ocean temperature; Martiny et al., 2013). Indeed, environmental variability in the supply of key elements like phosphorus (P) and nitrogen (N) is the single best predictor of differences in cellular concentrations of these elements among phytoplankton (Galbraith and Martiny, 2015). As well, species composition of local producer and consumer communities can influence carbon (C) and N concentrations in foliar tissues of plant species (Borer et al., 2015). This variability in elemental composition of autotrophs produces areas of high and low quality resources for herbivores across landscapes (Jean et al., 2015; Leroux et al., 2017). In turn, spatial heterogeneity in resource elemental composition – i.e., their stoichiometry (Sterner and Elser, 2002) – can influence consumers’ foraging strategies (Ball, Danell, and Sunesson, 2000; Youngentob et al., 2011). However, few studies to date have investigated how consumers’ space use varies in response to variability in resource stoichiometry (but see McNaughton et al., 1989). Here, we investigate how this mosaic of elemental hot and cold-spots in resource elemental composition (*sensu* Bernhardt et al., 2017; McClain et al., 2003) may influence the home range size of a small terrestrial mammal, the snowshoe hare (*Lepus americanus*).

The home range, the area an animal routinely uses to meet its daily needs (Burt, 1943; Powell and Mitchell, 2012), varies in size within and across species under the effect of multiple variables (Table 1; Tamburello, Côté, and Dulvy, 2015). For instance, the higher energetic needs arising from bigger body size lead to larger home range size (Peters, 1983). Likewise, diet composition and ecosystem function can also influence the size of an individual’s home range (Tamburello, Côté, and Dulvy, 2015). Carnivores tend to have larger home ranges than omnivores and herbivores due to the patchier nature of the resources they seek (Tamburello, Côté, and Dulvy, 2015; Tucker, Ord, and Rogers, 2014). As well, species living in low productivity habitats tend to have larger home ranges than species living in high productivity ones, because they need to move more to find enough food to avoid starvation (Tucker, Ord, and Rogers, 2014). For example, experimental evidence points to terrestrial herbivores responding to variability in both quantity and quality of their preferred resources at multiple spatio-temporal scales (e.g., Ball, Danell, and Sunesson, 2000; Nie et al., 2015). Yet, studies often rely on proxies to measure variability in resource quality and these proxies can vary among study systems, from forage species identity (van Beest et al., 2011), to nutritional value (e.g., carbohydrate content; Saïd et al., 2009), to forage availability (Duparc et al., 2020). Within the framework of ecological stoichiometry, resource quality is often defined based on the elemental composition of the resource — that is, its content of key nutrients such as C, N, and P (Leal, Seehausen, and Matthews, 2017). Here, we argue that spatial variability in a resource’s elemental composition may inform consumer space use.

**Table 1:** Environmental and ecological drivers of home range size among mammals.

| Variable | Effect | References |
| --- | --- | --- |
| Body Size | larger body mass corresponds to larger home ranges | Jetz (2004), Kelt and Van Vuren (2001), Mace and Harvey (1983), Ofstad et al. (2016), Peters (1983), and Tucker, Ord, and Rogers (2014) |
| Habitat | richer habitats usually corresponds to smaller home ranges | Dussault et al. (2005), Fridell and Litvaitis (1991), Ofstad et al. (2016), Tucker, Ord, and Rogers (2014), Walton et al. (2017), and Willems and Hill (2009) |
| Information | previous knowledge of an area's distribution of resources, risk sources, mates, and refugia varies how individuals use available space | Merkle et al. (2015), Powell and Mitchell (2012), Spencer (2012), and Zweifel-Schielly et al. (2009) |
| Diet | carnivores have larger home ranges than herbivores and omnivores | Kelt and Van Vuren (2001), Mace and Harvey (1983), and Tucker, Ord, and Rogers (2014) |
| Energy | increasing energetic demands lead to larger home ranges | Kelt and Van Vuren (2001) and Mcart et al. (2009) |
| Behaviour | sociality can reduce home range size by allowing for more efficient foraging (e.g., pack hunting) | Carbone, Teacher, and Rowcliffe (2007) |

Animals make space use decision at multiple spatio-temporal scales, from which food items to forage on in a patch to where to establish their geographic range(Johnson, 1980). Within a patch, available food items vary in their quality and quantity and animals, in turn, forage only on some of these food items (4^*th*^ order selection; Johnson, 1980). Koalas (*Phascolarctos cinereus*) and greater gliders (*Petauroides volans*) prioritize use of high-quality *Eucalyptus* spp. patches, albeit in different ways: while koalas search for and forage longer on trees whose leaves have high N concentrations (Marsh et al., 2014; Moore et al., 2010), greater gliders actively avoid those trees with high levels of N-based secondary metabolites in their leaves (Youngentob et al., 2011). Thus, animals tend to use some areas of the landscape more than others (3^*rd*^ order selection; Johnson, 1980). For instance, in Scandinavia, moose (*Alces alces*) and mountain hare (*Lepus timidus*) visited white birch (*Betula pubescens*) and Scots pine (*Pinus sylvestris*) more frequently in N-fertilized areas compared to unfertilized controls (Ball, Danell, and Sunesson, 2000). Home ranges arise from these patch use patterns. For example, bamboo-exclusive giant pandas (*Ailuropodia melanoleuca*) seasonally shift their range and vary their home range size in response to variation in N, P, and calcium content in their food — consistently foraging on the highest-quality food available as a result (Nie et al., 2015).

As these examples show, resource elemental composition can play an important role in determining how animals use their space: where they forage, what they forage on, for how long, and when. With the recent development of new statistical methods to predict resource stoichiometry at landscape extents (e.g., Galbraith and Martiny, 2015; Leroux et al., 2017; Soranno et al., 2019), we can investigate how resource elemental composition influences consumers’ distribution beyond the local patch. For example, stoichiometric distribution models (henceforth, StDMs) can predict element distributions over landscapes and allow identification of hot and cold spots of resource elemental composition across spatial extents (Leroux et al., 2017). StDMs allow for studying patterns of consumers’ space use and distribution in a stoichiometrically informed way. For instance, Leroux et al. (2017) used StDMs predictions to investigate the spatial distribution of moose (*A. alces*) at the landscape extent, in the boreal forests of northern Newfoundland. Spatial distribution models of moose performed consistently better when including a measure of forage elemental composition (e.g., elemental dry weight, % content, or ratios), providing evidence that spatial gradients in plant stoichiometry may influence herbivores’ space use decisions (Leroux et al., 2017). Consequently, as in the case of the giant panda mentioned above (Nie et al., 2015), spatial variability in forage elemental composition may also drive an animal’s home range size.

Here, we use elemental distribution models, i.e., StDMs, to investigate the relationship between summer home range size and resource elemental composition in snowshoe hares (*L. americanus*). Snowshoe hares are a keystone herbivore in the boreal forests of North America (Feldhamer, Thompson, and Chapman, 2003). Snowshoe hare habitat is strongly N and P-limited (Price et al., 2013), and these constraints influence their ecology, behavior, and physiology (Murray, 2002; Thornton et al., 2013). These characteristics make snowshoe hares uniquely suited to address these questions. We use stoichiometric ratios — e.g., C:N, C:P, N:P ratios — as proxies for resource quality for snowshoe hares. High C:N or C:P forage tends to be woody, hence less digestible, whereas high N:P forage may not offset the boreal forest’s strong P-limitation (Leroux et al., 2017; Townsend et al., 2007). Hence, we consider food items with low C:N, C:P, and N:P ratios as higher quality resources than those with high C:N, C:P, or N:P ratios. As well, snowshoe hares may respond to the overall quality of an area – i.e., the area’s average quality — or to the variation in quality within an area – i.e., how heterogeneous the quality of an area is (Zweifel-Schielly et al., 2009). Thus, we test the hypothesis that spatial differences in average resource quality, the variability of resource quality, or both influence snowshoe hare home range size (Figure 1). We predict that (i) snowshoe hares in areas of homogeneous resource quality (low variability) will have smaller home ranges than individuals in areas with more spatially heterogeneous resources. We further predict that (ii) snowshoe hares in areas of lower average forage C:N or C:P ratio will have smaller home ranges than individuals in areas in which these forage ratios are higher. For N:P ratio, we predict (iii) that snowshoe hares will have larger home ranges in areas of high N:P ratio, i.e. P-limited, than in areas of low N:P ratio, i.e. N-limited. Finally, we expect (iv) that in areas with low and spatially homogeneously stoichiometric ratios (low mean and low variation), snowshoe hares will have smaller home ranges compared to areas where these metrics are both high or where one is high and the other is low.

**Figure 1.**
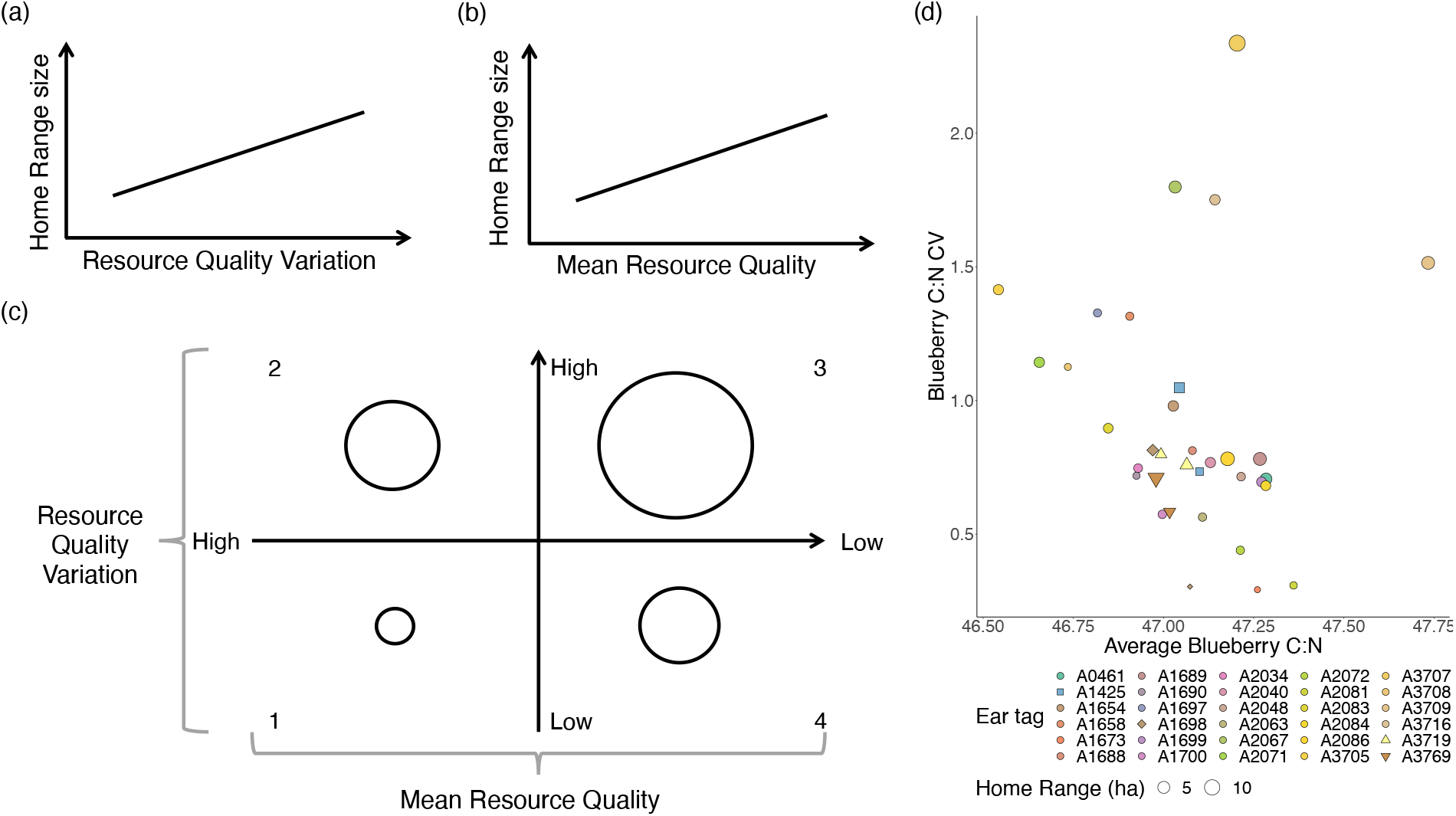
Predictions of the relationship between resource elemental composition and home range size in snowshoe hares. **(a)**: as variability in resource elemental composition increases, home range size will also increase, as per predictions (i) and (iii). **(b)**: with increasing average resource elemental composition, home range size will increase, as per predictions (ii) and (iii). **(c)**: jointly, these two dimensions of variation produce a “resource quality space”, where they interact to influence home range size — as per prediction (iv). In this space, where average resource quality is high and its variability low, herbivore home range size will be small (quadrant 1). Conversely, when variability is high and average quality is low, home range size will be large (quadrant 3). When either average quality is high and its variability is low or vice versa, home range size will be intermediate between the two extremes (quadrants 2 and 4). **(d)**: distribution of snowshoe hare home range size estimates (ha) from this study (*n* = 30) in lowland blueberry resource quality space, defined by foliage C:N ratio. Most hares in our sample live in areas of moderate lowland blueberry C:N content and variability. Some individuals maintain small home ranges in areas of relatively homogeneous, medium-to-high lowland blueberry foliage C:N ratio (e.g., A1673, A2702, A2081). Conversely, a few snowshoe hares with large home ranges live in areas of heterogeneous, low-quality lowland blueberry (e.g., A3705). The empty lower left corner may indicate that no areas of high and homogenous resource quality are available in our study area, or that no hares are using it if it is present. Data point size reflects 50% UD home range size; different colors identify different individuals. Different shapes separate individuals with more than one year of telemetry sampling (squares, A1425; diamonds, A1698; triangles, A3719; upside-down triangles, A3769) from individuals with only one year of telemetry (circles). The Supplementary Information contains additional details on the degree of overlap between home ranges from consecutive years for these four individuals (Table S3 and Figures S3 to S6).

## Methods

### Study Area

We conducted our study in four boreal forest stands in eastern Newfoundland, Canada, in and around Terra Nova National Park (48°31′50″ N, 53°55′41″ W;Figure S1). We selected forest stands based on snowshoe hare habitat preferences and along a forest stand age chronosequence with four categories; 20–40 years old, 41–60 y. o., 61–80 y. o., and 81–100 y. o. (see SI Section S3 for more details). In all four forest stands, black spruce (*Picea mariana*) dominates the canopy, which also comprises balsam fir (*Abies balsamea*), red maple (*Acer rubrum*), white birch (*Betula papyrifera*), and white spruce (*Picea glauca*). Lowland blueberry (*Vaccinium angustifolium*), Sheep laurel (*Kalmia angustifolia*), and Labrador tea (*Rhododendrum groenlandicum*) dominate the understory. In May 2016, we established a 500m×500m snowshoe hare live trapping grid housing 50 Tomahawak live traps (Tomahawk Live Trap Company, Hazelhurst, WI) along a meandering transect in each of the four forest stands (see SI section S3 and fig. S2).

### Spatial Variability in Food Stoichiometry

We collected plant samples, ~20 g wet weight, in and around each trap location on the four live-trapping grids during the summer months of 2016 and 2017. We focused on three important summer forage species for snowshoe hares (Dodds, 1960): lowland blueberry (*V. angustifolium*), red maple (*A. rubrum*), and white birch (*B. papyrifera*). Our sampling attempted to replicate hare browsing by collecting only new growth material — that is, new leaves and terminal ends of branches. We shipped 10 g dry weight from each sample to the Agriculture and Food Laboratory at the University of Guelph to measure content of C, N, and P for each of our three plant species of interest (listed above; henceforth, SOI).

In our analyses, we used quantitative predictions of foliar C:N, C:P, and N:P ratios obtained from fitting StDMs to the stoichiometry data obtained from plant samples from all four grids. We built five StDMs. Here we briefly describe the procedure behind building and fitting StDMs (see Heckford et al., n.d., *in revision* for detailed methods and Leroux et al., 2017, for general background on StDMs). To build each StDM, we used three types of plant SOI data: (i) sampling plot density data from a shrub belt sampled along the South-North diameter (22.6m) of the plot, divided into 4 height classes; (ii) elemental percentages, i.e., % C, N, P, extracted from foliar samples; and (iii) biomass data collected in areas adjacent to our sampling grid. We first fit allometric models for each study species using the formula: *log*(*biomass*) ~ *log*(*basal diameter* + *height*). At the sampling plot level, this allowed us to estimate density of plant SOI by height class based on shrub belt data, and to use these estimates to predict plant SOI biomass by height class in each sampling plot. We then calculated C, N, P foliar content per SOI per plot by dividing a SOI’s total plot biomass by the product of plot area and foliar elemental content (% dry weight). We obtained C, N, P quantity estimates by dividing elements’ foliar content by their molar weight, and stoichiometric ratios from these estimates (C:N, C:P, N:P; Heckford et al., n.d., *in revision*).

Each StDM included spatially explicit covariates, grouped into four categories: land cover, productivity, biotic, and abiotic factors. Preliminary analyses of yearly variation in plant SOI stoichiometry showed negligible variability between 2016 and 2017 (Richmond et al., n.d., *in review*). Hence, we did not include year of sampling as a covariate in our StDMs. We fit a set of 15 Generalized Linear Models based on *a priori* hypotheses (see Heckford et al., n.d., *in revision*), including a null model, to nine response variables: percent element content (% C, N, P), quantity element content (C, N, P, *g*/*m*^2^), and stoichiometric ratios (C:N, C:P, N:P). We used the Akaike Information Criterion corrected for small sample size (AICc; Burnham and Anderson, 2002) to assess the weight of evidence supporting each model. After removing uninformative parameters (*sensu* Leroux, 2019), we used the top-ranked model for each SOI-stoichiometric ratio pair to produce predictive plant SOI stoichiometry surfaces as proxies for resource quality available within hare home ranges.

### Home range size and stoichiometry

In May-November of 2016 through 2019, we live-trapped and radio-collared snowshoe hares in the youngest forest stands, 20–40 years old (henceforth, hare study area). We baited each trap at dusk with apple slices, alfalfa, and rabbit chow, and checked them the following dawn. We collected body weight (g) and other demographic data of each hare, before fitting it with a 25 g radio collar (M1555, Advanced Telemetry Systems, Isanti, MN) and releasing it. We did not fit individuals with radio-collars when the weight of the collar was ≥5% of the hare’s own body weight. The Animal Care Committee of Memorial University of Newfoundland approved our live-trapping and handling protocol with permit 18-02-EV. Further details on our live-trapping protocol can be found in SI section S3.

In May-September of 2017 through 2019, we located snowshoe hares using handheld receivers (Biotracker, Lotek, Ontario, CA) and VHF antennas (RA–23K, Telonics, AZ). We collected three or more azimuths per hare per day, storing them in an electronic data collection form on an iPad (FileMaker Pro Advanced, v. 14; Claris International Inc., 2015) and using digital maps (Avenza Maps, v. 3.7; Avenza Systems Inc., 2020) to check the triangulation polygon’s size. We estimated home range size and ran all subsequent analyses in R (v. 4.0.1; R Core Team, 2020). For each hare in our sample (*n* = 30), we used package `razimuth` to estimate collar location based on the Azimuthal Telemetry Model (Gerber et al., 2018). From these locations, we estimated the Utilization Distribution (UD) of our snowshoe hares using the autocorrelated Kernel Density Estimator corrected for small sample size (AKDEc) using the `ctmm` R package (Fleming and Calabrese, 2017; Fleming, Noonan, et al., 2019). From the UDs, we estimated home range area in hectares (ha) at the 50%, 75%, and 90% isopleths. For more details on our home range estimation workflow, please see SI section S5 and the Supporting Code document.

We used function `extract` from the `raster` R package (Hijmans, 2020) to overlay the boundary of each snowshoe hare’s home range area estimate, i.e, the 50%, 75%, 90% UD isopleths, on the stoichiometric surfaces and get C:N, C:P, and N:P values for every pixel covered by the home range (see Supporting Code for more details). From these data, for each home range, we estimated (i) each stoichiometric ratio’s mean value and (ii) its coefficient of variation. The coefficient of variation (henceforth, CV), the ratio of a sample’s standard deviation to its mean value, provides an easy-to-interpret assessment of how variable the predicted SOI stoichiometry of a given home range is, compared to its mean value. See Supporting Code for more details.

### Statistical Analyses

We used linear models to investigate the effects of resource stoichiometry, i.e., mean, CV, and their interactive effects, and body weight on the size of the home range of snowshoe hares estimated at the 50% (i.e., the core area; Börger et al., 2006), 75%, and 90% isopleths. We included body weight to capture potential intraspecific variability in home range size due to an individual’s ecology and physiology (Peters, 1983). Conversely, we did not include year of sampling, as preliminary analyses provided no evidence it influenced home range size of our snowshoe hares (see Supplementary Code; Börger et al., 2006). As well, we did not include sex in our models as evidence for snowshoe hares points to this variable being correlated with body weight (Feldhamer, Thompson, and Chapman, 2003) and does not appear to influence the elemental composition of snowshoe hares (Rizzuto et al., 2019). For each combination of plant SOI and C:N, C:P, and N:P (*n* = 5), to test prediction (i) we fit a model including each stoichiometric ratio’s CV. Likewise, to test predictions (ii) and (iii) we fit a model including the ratios’ mean values. To test prediction (iv) we fit a model including the additive effects and a model including the additive and interactive effects of the ratios’ mean and coefficient of variation. For each model, we also fit a version that included the hares’ body weight. We fit this set of 8 models, plus a null model, to our dataset and used function `AICc` in the `AICcmodavg` R package to select top models based on parsimony (Burnham and Anderson, 2002; Mazerolle, 2017). Following Leroux (2019), we removed uninformative parameters from the model set of each stoichiometric ratio. Below, we report summary AICc tables and refer the interested reader to the Supporting Code document for full AICc tables.

## Results

StDMs of red maple C:N, N:P ratios, and lowland blueberry C:N, C:P, N:P ratios all ranked above the null model whereas all other StDMs (i.e., red maple C:P ratio, white birch C:N, C:P, N:P ratios) were not supported by the data (Heckford et al., n.d., *in revision*). We used this suite of five StDMs to produce geo-referenced predictions of resources’ spatial variability in and around our hare study area.

Our sample of radiocollared snowshoe hares included 30 individuals: 4 followed during summer 2017, 6 in summer 2018, and 20 during summer 2019. We followed four snowshoe hares for two consecutive sampling years: three in the 2018 and 2019 sampling seasons and one in the 2017 and 2018 sampling seasons. For the individuals sampled in more than one year, we included in the analyses only the home range size estimate from the year with the most telemetry points. Our results are not sensitive to this decision (see Supplementary Code). Our sample included 14 females, 12 males, and 4 individuals of unknown sex. Adult hares comprised the majority of our sample (n=27), with two young-of-the-year and one unknown. Mean core area size was 4.292 ha (range: 0.835–11.465) for 2017, 3.104 ha (range: 0.215–6.163) for 2018, and 2.68 ha (range: 0.486–7.403) for 2019 (3-year mean± SD: 2.996 ha ± 2.300). For lowland blueberry, within the core area, predicted C:N ratio ranged from 45.32 to 49.17 (median: 47.15), predicted C:P ratio ranged from 1201 to 2277 (median: 1279), and predicted N:P ratio from 25.15 to 45.42 (median: 28.09). For red maple, predicted C:N ratio ranged from 23.26 to 39.79 (median: 30.89) and predicted N:P ratio ranged from 28.39 to 39.09 (median: 34.13).

We found mixed support for prediction (i), resource quality heterogeneity influencing home range size. The CV of lowland blueberry C:N ratio and red maple N:P ratio appeared in the top models for home range core area size (slope = 3.429 ± 0.664, *R*^2^ = 0.548, and slope = 0.866 ± 0.378, *R*^2^ = 0.15, respectively; Table 2 and fig. 2). This trend holds at all kUD isopleths for lowland blueberry, but not for red maple (Tables S1 and S2). Indeed, the CV of lowland blueberry C:N ratio explained a higher portion of the variation in snowshoe hare home range size, compared to the mean value of this ratio (CV-only model *R*^2^ = 0.376, mean-only model *R*^2^ = 0.102; Table 2). We found no evidence of this relationship for the CV of lowland blueberry C:P, N:P ratios, and only weak evidence supporting this trend for the CV of red maple C:N ratio for home range size estimates at 50% (slope = 0.127 ± 0.089, *R*^2^ = 0.166; Table 2) and 75% kUD (Table S1).

**Table 2:** Top ranking GLMs describing the relationship between home range core area and resource stoichiometry, after removing uninformative parameters (see Supporting Code for full AICc tables). For each plant SOI and stoichiometric ratio pair, we report the top model, any model above the intercept, and the intercept. For coefficients, we report values as *estimate* (±*SE*). Column headers: K, number of parameters in the model; LL, log-likelihood; CV, Coefficient of Variation; BW, body weight.

|  |  |  |  | Coefficients |  |  |  |
| --- | --- | --- | --- | --- | --- | --- | --- |
| K | $\Delta AIC_c$ | LL | R<sup>2</sup> | Intercept | Mean | CV | BW |
| Blueberry C:N top models |  |  |  |  |  |  |  |
| 4 | 0.000 | −55.783 | 0.548 | -199.097<br>(±62.029) | 4.224<br>(±1.316) | 3.419<br>(±0.664) |  |
| 3 | 7.025 | −60.634 | 0.376 | 0.058<br>(±0.789) |  | 3.110<br>(±0.757) |  |
| 3 | 17.952 | −66.098 | 0.102 | -148.049<br>(±84.809) | 3.208<br>(±1.802) |  |  |
| 2 | 18.792 | −67.707 | 0.000 | 2.974<br>(±0.430) |  |  |  |
| Blueberry N:P top models |  |  |  |  |  |  |  |
| 3 | 0.000 | −65.859 | 0.116 | -3.230<br>(±7.684) | 0.327<br>(±0.245) |  | -0.002<br>(±0.001) |
| 2 | 0.033 | −67.707 | 0.000 | 2.974<br>(±0.430) |  |  |  |
| Blueberry C:P top models |  |  |  |  |  |  |  |
| 3 | 0.000 | −65.859 | 0.116 | -7.999<br>(±5.741) | 0.008<br>(±0.004) |  |  |
| 2 | 1.218 | −67.707 | 0.000 | 2.974<br>(±0.430) |  |  |  |
| Red Maple N:P top models |  |  |  |  |  |  |  |
| 3 | 0.000 | −65.130 | 0.158 | 0.162<br>(±1.291) |  | 0.866<br>(±0.378) |  |
| 2 | 2.676 | −67.707 | 0.000 | 2.974<br>(±0.430) |  |  |  |
| Red Maple C:N top models |  |  |  |  |  |  |  |
| 4 | 0.000 | −64.980 | 0.166 | 5.080<br>(±2.338) |  | 0.127<br>(±0.089) | -0.002<br>(±0.001) |
| 3 | 0.000 | −66.318 | 0.088 | 1.644<br>(±0.908) |  | 0.149<br>(±0.090) |  |
| 2 | 0.299 | −67.707 | 0.000 | 2.974<br>(±0.430) |  |  |  |

**Figure 2.**
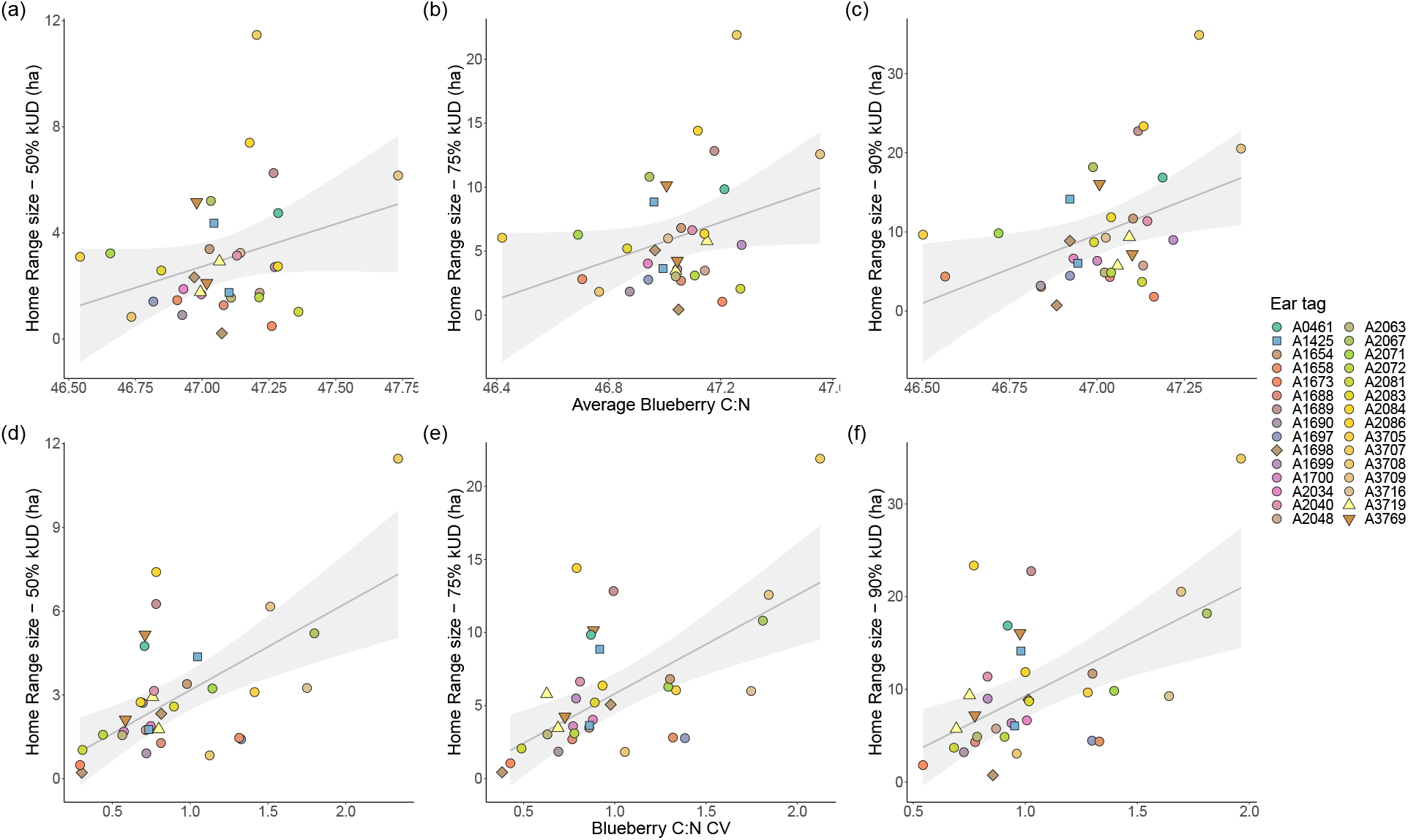
Relationship between lowland blueberry foliage C:N ratio quality metrics and home range size, at 50%, 75%, and 90% UD. **Upper panels**: the size of home range core area for our snowshoe hares is smaller in areas of lower mean lowland blueberry foliage C:N ratio and increases with the ratio’s mean value (panel a). Home range sizes estimated from larger isopleths shows similar trends (panels b, c). Higher values of C:N ratio point towards lower availability of N in blueberries, so that individuals living in such areas (e.g., A2084) may have to forage over larger areas to meet their elemental requirements of N to survive (Sterner and Elser, 2002). **Lower panels**: at increasing values of the variability in lowland blueberry foliage C:N ratio corresponds a sharp increasing in home range core area size of snowshoe hares (panel d), a trend repeated at larger isopleths (panels e, f). Snowshoe hares in areas of high variability of lowland blueberry N content may resort to foraging over much larger areas than individuals that have access to food items of less variable quality — regardless of whether this is high or low quality. Grey lines are regression lines drawn from the top-ranking model for lowland blueberry C:N ratio at the relevant UD isopleth (Table 2 and Tables S1 and S2) and light grey shaded areas around them represent 95% Confidence Intervals. All other specifications as in Figure 1.

We found mixed support for our prediction (ii) on the effects of average C:N ratio and C:P ratio on home range size. The mean values for lowland blueberry foliage C:N ratio and C:P ratio appeared in top models (slope = 4.224 ± 1.316, *R*^2^ = 0.548, and slope = 0.008 ± 0.004, *R*^2^ = 0.116, respectively; Table 2 and fig. 2), with the trend holding at all three isopleths for average C:N ratio, but only at the 75% isopleth for average C:P ratio (Tables S1 and S2). While the top model included both mean and CV of lowland blueberry C:N ratio, the mean-only model was ranked 3rd overall and explained 10% of the variation in hare home range size (Table 2). No support for this prediction came from models using average red maple C:N ratio (Tables S1, 2 and S2). As well, we found weak evidence supporting prediction (iii), home range size increasing as resources’ N:P ratio increases, from lowland blueberry foliage (slope = 0.327 ± 0.245, *R*^2^ = 0.159; Table 2) but the trend does not hold at either the 75% or 90% isopleth (Tables S1 and S2). We found no support for prediction (iv) at any kUD isopleth for C:N and C:Pany of the three ratios considered, as the top models for lowland blueberry C:N ratio and C:P included the additive effects of mean and CV and only the mean, respectively (Table 2, and Tables S1 and S2).

## Discussion

Animals forage on a variety of resources whose elemental composition may influence space use and foraging patterns at multiple spatial scales (Duparc et al., 2020; Lima and Zollner, 1996; van Beest et al., 2011). We found evidence that spatial differences in a preferred resource’s predicted average elemental composition or its variability correlated with herbivore home range size. Additionally, forage species identity may also play a role, further influencing these relationships. Together, our results provide evidence supporting the role that resources’ elemental content plays in influencing consumers’ spatial ecology. Our results suggest that exploring the fundamental question of animal space use through an elemental lens may allow researchers to better trace the feedbacks between animals and ecosystem functions, e.g., elemental cycling (Schmitz et al., 2018).

The boreal forest is a strongly N and P-limited ecosystem (Price et al., 2013). Snowshoe hares need to carefully balance their intake of C-heavy plant food against their N and P growth requirements (Sterner and Elser, 2002). Our results provide explicit evidence of this elemental trade-off at the home range scale and highlight how differences in resource elemental phenotype within and across areas used by snowshoe hares underlie variation in home range size in a heterogeneous landsacpe. In particular, results for both lowland blueberry foliage C:N ratio and red maple foliage N:P ratio support prediction (i), that variability in N and P content within a home range core area can influence its size (Table 2). Snowshoe hares in our study appear to readily respond to stoichiometric changes in lowland blueberry, one of their preferred summer forage (Dodds, 1960). Variability in the elemental phenotype of the main components of a consumer’s diet appears to influence both their spatial and temporal distribution over the environment (McNaughton et al., 1989; Nie et al., 2015). In our study area, lowland blueberry is more abundant than red maple as well as, overall, more browsed (SI Figure S7). A higher sensitivity to the variability in quality of this resource, then, may point to the elemental composition of these two plant species playing a fundamental role in a snowshoe hare’s efforts to meet its high nutritional requirements (Murray, 2002).

Furthermore, we find evidence that elements can influence home range core area size even when considering an area’s average quality — i.e., when smoothing the variation to a single value — in accordance with predictions (ii) and (iii). In particular, low average values of C:N, C:P, and N:P ratios for lowland blueberry consistenly correspond to smaller home range size (Table 2 and Figure 2). This held true for C:N ratio whether estimated home range size from the core area or from larger UD slices — suggesting that resource quality may influence space use decisions at a higher order of selection (i.e., landscape or 3^*rd*^ order of selection Johnson, 1980). Interestingly, we additionally find evidence that a ratio’s coefficient of variation may add an additional side to this relationship, as it appears in the top model for lowland blueberry foliage C:N ratio at all three UD slices. Indeed, hares living in areas of high mean and high coefficient of variation for the foliage C:N ratio of lowland blueberry appear to have larger home ranges than those living in areas where mean values are high but the coefficient of variation is small (Figure 1, panel d, Table 2, and Tables S1 and S2). Thus, consumers may use different information cues to make space use decisions at different spatial scales — e.g., acros vs. within patches on the landscape.

Similar effects of resource quality on herbivore space use patterns have been described in other study systems. For instance, other species of leporids, as well as ungulates, tend to increase use of areas where they have access to forage with higher content of limiting nutrients (Ball, Danell, and Sunesson, 2000). In turn, this preferential use of areas where forage is high in limiting nutrient content appears related to reproductive and physiological benefits (Mcart et al., 2009) or to population dynamics (Merkle et al., 2015). Overall, the elemental composition of forage items appears to be a fundamental driver of herbivore space use across spatial scales; from which food items to eat within a patch, to which habitats to establish a home range in, to which areas to visit over the landscape (Ball, Danell, and Sunesson, 2000; Nie et al., 2015; Zweifel-Schielly et al., 2009).

Evidence of resource quality influence on space use decisions of consumers arising from several study system corroborates this result (e.g., Nie et al., 2015; Saïd et al., 2009; van Beest et al., 2011). Indeed, the majority of the hares in this study appear to live in areas of relatively high N and P values in the foliage of both red maple and lowland blueberry (Figure 1, panel d). The few cases of use of areas with high resource heterogeneity may result from population dynamics, particularly the increase in hare numbers from 2017 to 2019. In 2017, our collared snowshoe hares all had home ranges in relatively high quality areas for lowland blueberry. As more snowshoe hares appeared on the landscape in 2018 and 2019, new individuals increasingly established larger home ranges that extended beyond the areas of lower heterogeneity or higher overall N or P availability. Furthermore, the high degree of overlap we found between home range estimates calculated for hares with more than one year of telemetry data may point to a limited ability of older snowshoe hares to retain their range across years (Table S3 and Figures S3 to S6). Other herbivores appear to have similar growth-dependent colonization of less-favorable areas of a landscape. Among bison (*Bison bison*), individuals appeared to expand their population range to include areas of lower resource quality and establish larger home ranges in them as population density increased over time (Merkle et al., 2015). Similar patterns of population spatial distribution driven by resource availability and foraging strategies are fairly well-known among passerine birds (Piper, 2011). The elemental composition of foraging resources, then, may not only influence the size of a consumer’s home range, but also its location over the landscape. However, to our knowledge, this study is the first to show that key chemical elements may drive animal space use decisions.

We modeled our measure of forage quality, forage stoichiometry, based on a suite of environmental, biotic, and abiotic covariates (Heckford et al., n.d., *in revision*). This approach may help investigate direct drivers of consumer space use and shed light on ecosystem characteristics allowing high-quality resources to persist in an area. In turn, the environmental drivers that correlate with forage stoichiometry may indirectly influence a consumer’s spatial ecology even in the absence of the resource itself. Further, StDMs allow accounting for multiple ecological currencies shaping a consumer’s ecology at varying spatio-temporal scales (Levin, 1992; Lima and Zollner, 1996). Thus, applying stoichiometric measures of forage to model consumer space use may be a fundamental tool in bridging metabolic, nutritional, landscape, and behavioural ecology (Sterner, 2004). In turn, this may allow us to disentangle the ubiquitous relationships and feedbacks among consumer, resources, and the environmental and ecological processes they are part of (Levin, 1992; Lima and Zollner, 1996). Furthermore, our StDM-driven approach explains a large portion of the variance observed in our sample, albeit with some variability among model sets (see Tables S1 and S2). Indeed, the elemental composition of resources has been shown to accurately describe and predict the spatial distribution patterns of consumers in a variety of biomes, from boreal (this study), to tropical (McNaughton et al., 1989), to temperate (Merems et al., 2020; Nie et al., 2015).

Overall, our results provide evidence that ecological stoichiometry may help researchers understand fundamental components of consumers’ space use. Based on the emergent field of spatial stoichiometry (Galbraith and Martiny, 2015; Leroux et al., 2017; Soranno et al., 2019) and our own results, we argue that using the elemental composition of resources to investigate patterns of consumer space use may provide a comparable and potentially more parsimonious approach than other, more widespread methods — e.g., habitat classification (Zweifel-Schielly et al., 2009), forage species identity (van Beest et al., 2011), or availability (Duparc et al., 2020). Focusing on stoichiometric currencies would allow for consistency in defining and measuring fundamental metrics, e.g., resource quality, across studies and study systems. It would also reduce the need to rely on elemental conversion factors, increasingly recognized as problematic due to their lack of generality across different food items and outdated estimation methods (Mariotti, Tomé, and Mirand, 2008). As well, stoichiometric currencies may help investigate the different experiential layers that make up an animal’s home range (*sensu* Powell and Mitchell, 2012), further refining how researchers measure, describe, and interpret animal space use at multiple spatio-temporal scales (Levin, 1992). Finally, rooting theoretical models of ecological processes in stoichiometric units may make them more widely applicable to real world scenarios (Schmitz et al., 2018).

Life builds itself using a limited subset of elements (Kaspari and Powers, 2016). These are continuously transformed and exchanged, globally, among organisms and their abiotic environment, and within and across ecosystem borders. Ecological stoichiometry offers an ultimately reductionist approach that, by providing common units of measurement with which to describe both actors and currencies involved in these exchanges, may effectively provide researchers with a holistic perspective to explore animal space use.

## Data Availability

All the data and code used in the analyses are available in a figshare repository: https://doi.org/10.6084/m9.figshare.12798296.v1.

## Acknowledgements

We thank J. Strong, A. Hann, B. Stratton, K. Gerrow, and G. R. Riesefel for their help during data collection. We thank J. Kennah for helpful comments on earlier drafts of the manuscript. We thank Dr. S. Berg, Dr. C. Fleming, and S. Andrews for help with sigloc, ctmm, and spatil R, respectively. This research was funded by the Government of Newfoundland and Labrador Centre for Forest Science and Innovation (to SJL, YW, and EVW), Government of Newfoundland and Labrador Innovate NL Leverage R&D (to EVW & SJL) and Ignite R&D (to SJL) programs, Mitacs Accelerate Graduate Research Internship program (to YW, EVW, & SJL), the Canada Foundation for Innovation John R. Evans Leaders Fund (to EVW & SJL), and a Natural Science and Engineering Research Council Discovery Grant (to SJL). M. Rizzuto, S. J. Leroux, Y. F. Wiersma, and E. Vander Wal designed the study; M. Rizzuto, T. Heckford, J. Balluffi-Fry, I. C. Richmond, Y. Wiersma, and S. J. Leroux collected the data; M. Rizzuto, S. J. Leroux, and I. C. Richmond analyzed the data. All authors contributed to interpreting the results. M. Rizzuto led the writing of the manuscript and all authors read and approved the final version.

