## Supplementary Information for "Forage stoichiometry predicts the home range size of a small terrestrial herbivore"

Version dated: August 13, 2020

#### Contents

**S1 Introduction** **2**

**S2 Study species** **2**

**S3 Grid Establishment and Live Trapping** **3**

**S4 Triangulation and Data Processing** **4**

**S5 Home Range Estimation** **4**

S5.1 Accounting for pseudoreplication . . . . 4

S5.2 Testing for Sampling Year-related effects . . . . 5

**S6 Supplementary Tables** **5**

**S7 Supplementary Figures** **9**

### S1 Introduction

The present document contains supplementary information on the study species, area, and methods used in our paper investigating the influence of forage elemental composition on the home range size of snowshoe hares (*Lepus americanus*) in the boreal forest of the island of Newfoundland. Section S2 provides additional details on the ecology of the snowshoe hare, a keystone herbivore species in the boreal ecosystem. Section S3 describes the rationale and techniques used in establishing our snowshoe hares live-trapping grids, and Section S4 provides details on our triangulation protocol. We provide details on home range size estimation in Section S5, describing our handling of pseudoreplicates in Section S5.1. In Section S6, we provide supplementary tables including ΔAICc-ranked model selection tables for the analyses ran on home range size estimated at the 75% and 90% UD isopleths. Finally, in Section S7, we provide visual supporting materials, including a map of our study area, the layout of our live-trapping grids, and maps showing the degree of overlap among home range estimates from consecutive sampling year for four snowshoe hares.

We refer the interested reader to the companion R notebook "Supporting Code" for an in-depth description of the R workflow and code used to perform UD home range size estimation, overlap analyses, stoichiometric data extraction, and model fitting, as well as additional visual supporting materials including Utilization Distribution maps and an interactive map of our study area.

### S2 Study species

The snowshoe hare (*L. americanus*) is a keystone herbivore in the boreal forest and has been extensively studied for the role its characteristic boom-bust population cycle plays in the boreal forest's ecosystem dynamic (Krebs, Boonstra, and Boutin, 2018). Snowshoe hare populations cycle regularly in their abundance, with an 8–11 years periodicity that was already evident to $19^{th}$ century fur trappers (Feldhamer, Thompson, and Chapman, 2003; Krebs, Boonstra, and Boutin, 2018). Active year-round, snowshoe hares (*L. americanus*) seasonally forage on different plant species (see Dodds, 1960, for an extensive list) and face consistently high predation risk from a diverse group of both land-based and airborne predators (Krebs, Boonstra, and Boutin, 2018). Among these predators, the lynx (*Lynx canadensis*) is the most well-known, as its own population dynamics closely follow that of the snowshoe hare (Krebs, Boonstra, and Boutin, 2018). Furthermore, snowshoe hares have low fat tissue accumulation, $\leq$ 5% of their body weight on average, and vary their diet seasonally (Feldhamer, Thompson, and Chapman, 2003). They face high reproductive investment, with up to four litters per year and 6–8 leverets per litter (Feldhamer,

Thompson, and Chapman, 2003). Thus, access to good quality resources is paramount for their survival throughout the winter and to support their reproductive costs in spring and summer (Murray, 2002). Finally, in south-eastern portions of their range — like the island of Newfoundland — their distribution is patchier and more discontinuous compared to core areas, like the Yukon (Thornton et al., 2013; Krebs, Boonstra, and Boutin, 2018). As well, recent evidence points to their population cycle being less regular and shorter in Newfoundland than in the core areas of their range, like the Yukon (e.g., 8–9 years Reynolds et al., 2017)re In turn, this makes hares likely more sensitive to differences in resource quality and its variation over the landscape (Thornton et al., 2013).

#### S3 Grid Establishment and Live Trapping

We selected forest stands to house live-trapping grid using data from the Forest Resource Inventory compiled by the Provincial Government of Newfoundland and Parks Canada. We selected two forest stands inside the border of Terra Nova National Park, and two forest stands outside the protected area (Figure S1). We selected forest stands based on snowshoe hare habitat preferences and along a stand age chronosequence, with age groups 20–40, 41–60, 61-80, 81–100 years old. To reduce error in deploying the traps, we drew each grid's trap lines layout in ArcGIS (v. 10.4, ESRI, Redlands, CA) and then used digital, geo-referenced maps showing the location of each trap to deploy them. Upon deployment, we further confirmed trap location by taking averaged waypoints with a GPS unit (GPSMAP 64s, Garmin Ltd., Olathe, KS).

Live-trapping sessions lasted 3–4 days, with a pattern of one night of trapping, followed by one night of no trapping, followed by 1–2 additional trapping nights. In 2016 and 2017, we also used an optional pre-baiting night — i.e., baiting with the trap door jammed open — to allow individuals to familiarize themselves with the traps. Live-trapping nights depended on weather conditions: as snowshoe hares rely on movement to thermoregulate, due to their low body fat reserve (Feldhamer, Thompson, and Chapman, 2003), trapping in cold or wet nights may jeopardize their survival. Hence, whenever weather forecasts would call for night temperatures below 5 °C or for intense precipitation, e.g., rain or snow, we would suspend live-trapping. All live trapping followed animal care protocol 18-02-EV, approved by Memorial University's institutional Animal Care committee.

Before handling begun, we transferred trapped individuals from the trap to a burlap sack of known weight (~200 g). We inserted the door-end of the trap in the burlap sack, then opened the door and waited for the hare to walk in the sack. Once the hare was in the sack, we weighted (g) the whole sack. We then eartagged the individual and fitted it with a radio-collar before proceeding to collect data on sex, age, left hind foot length (mm), percent white fur estimated visually, and tick load. After releasing the individual, we calculated net wet body weight of the trapped hare by subtracting the weight of the empty burlap sack from the weight of the burlap sack with the hare in it. We further weighted the burlap sack between handling different individuals, to account for increased weight due to the sack getting soaked in rain and/or morning dew.

During our initial trapping season of 2016–2017, we had low capture rates and most of the individuals trapped and fitted with radiocollars were on the live-trapping grid located in the 20–40 years old forest stand (cfr., "hare study area" in the main text). Because of this, we decided to focus our trapping efforts on this grid and trapped in both spring and fall for the subsequent three years.

### S4 Triangulation and Data Processing

Triangulation happened daily from May to September in 2017, 2018, and 2019. We attempted to triangulate all active collars present on the grid every day: to avoid pseudoreplication, we randomized the order and time at which we attempted to locate each collar. In summer 2017, a single operator (MR) performed all triangulations, whereas in summers 2018 and 2019 three operators collected azimuths simultaneously from different locations: this greatly reduced the time necessary to triangulate all individuals and improved precision of triangulations. We estimated triangulation error for two of our operators using the `razimuth` R package (Gerber et al., 2018). In 2017, our sole observed had a median (±SD) error of 148 (±88) m, whereas in 2018 one out of two of our observers had a mean error of 265 (±370) m.

In R, we used package `razimuth` (Gerber et al., 2018) to estimate the location of each snowshoe hare for each set of azimuths collected daily during the sampling season. `razimuth` fits an Azimuthal Telemetry Model (ATM) to telemetry data using a Markov Chain Monte Carlo process. The ATM estimates both the location of the transmitter, in our case a radio-collar, and an error ellipses encircling it, which represents the uncertainty around the transmitter location (Gerber et al., 2018). These error ellipses can then be used to inform home range estimation using the ctmm package (Fleming and Calabrese, 2017; Fleming, Noonan, et al., 2019).

### S5 Home Range Estimation

We used an autocorrelated kernel density estimator (henceforth, aKDE) method to estimate home range area in hectares (ha) of our snowshoe hares, as this method is more reliable and accurate than more traditional approaches even with low relocation sample size (Fleming and Calabrese, 2017; Fleming, Noonan, et al., 2019). Below we describe how we addressed the few pseudoreplication issues that arose in our sample. The companion document "Supporting Code" contains code and maps for our home range size estimation.

#### S5.1 Accounting for pseudoreplication

For four snowshoe hares in our sample, we had telemetry data for two consecutive sampling years: three individuals were collared in the 2018 sampling season and were followed again in the 2019 sampling season, and one individual was collared in the 2017 and was followed again in the 2018 sampling seasons. We used the Bhattacharyya's affinity index (Fieberg and Kochanny, 2006) to estimate overlap in the utilization distribution of these four individuals between consecutive years (Winner et al., 2018). Results from this analysis highlight a high degree of overlap, likely an indication of strong site-fidelity across years Table S3. Armed with this knowledge, we addressed this issue in two ways: we decided to use only one year of sampling for each of these four individuals, and we used only the year with the most locations for each individual.

##### S5.2 Testing for Sampling Year-related effects

Evidence in the literature suggests that year of sampling — that is, the year during which relocations are collected — can play a role in determining home range size (Börger et al., 2006). Because of this, we ran preliminary analyses on our data using a model that included only the variable "sampling year", including only the year with most locations for the four individuals with year replicates (see section Section S5.1). We found no evidence of a relationship between sampling year and home range size (see companion document "Supporting Code" for code and model output). Alternative methods of considering the four individuals with year replicates, e.g., randomization, selecting the year closest to plant SOI sampling, yielded qualitatively similar results. Based on these results, we decided to not include the variable sampling year in our analyses.

#### S6 Supplementary Tables

Here we provide additional tables from our analyses of the influence of forage elemental composition on home range size of snowshoe hare. Table S1 and Table S2 show top ranking models for the analyses ran on home range size estimates at 75% and 90% aKDE isopleths, respectively. For full AICc tables, we refer readers to the companion document "Supporting Code". Table S3 provides details on the degree of overlap between home range estimates for the four individuals for which we had two years of telemetry data.

**Table S1:** Top ranking GLMs describing relationship between home range area at the 75% aKDE isopleth and resource stoichiometry, after removing uninformative parameters (see Supporting Code for full AICc tables). For each plant SOI and stoichiometric ratio pair, we report the top model, any model above the intercept, and the intercept. For coefficients, we report values as *estimate* ($\pm SE$). Column headers: K, number of parameters in the model; LL, log-likelihood; CV, Coefficient of Variation.

| | | | | **Coefficients** | | |
|---|---|---|---|---|---|---|
| K | ΔAICc | LL | $R^2$ | Intercept | Mean | CV |
| **Blueberry C:N top models** | | | | | | |
| 4 | 0.000 | −76.391 | 0.530 | -391.257 (±133.419) | 8.293 (±2.835) | 7.003 (±1.448) |
| 3 | 5.583 | −80.522 | 0.381 | -0.964 (±1.804) | | 6.770 (±1.629) |
| 3 | 16.047 | −85.754 | 0.123 | -348.474 (±178.606) | 7.537 (±3.797) | |
| 2 | 17.517 | −87.728 | 0.000 | 5.996 (±0.837) | | |
| **Blueberry N:P top models** | | | | | | |
| 2 | 0.000 | −87.728 | 0.000 | 5.996 (±0.837) | | |
| **Blueberry C:P top models** | | | | | | |
| 3 | 0.000 | −86.128 | 0.101 | -15.218 (±11.978) | 0.015 (±0.008) | |
| 2 | 0.721 | −87.728 | 0.000 | 5.996 (±0.837) | | |
| **Red Maple N:P top models** | | | | | | |
| 2 | 0.000 | −87.728 | 0.000 | 5.996 (±0.837) | | |
| **Red Maple C:N top models** | | | | | | |
| 3 | 0.000 | −86.318 | 0.090 | 2.820(±2.078) | | 0.323 (±0.194) |
| 2 | 0.341 | −87.728 | 0.000 | 5.996 (±0.837) | | |

**Table S2:** Top ranking GLMs describing relationship between home range area at the 90% aKDE isopleth and resource stoichiometry, after removing uninformative parameters (see Supporting Code for full AICc tables). All specification as in Table S1.

| | | | | **Coefficients** | | |
|---|---|---|---|---|---|---|
| K | ΔAICc | LL | $R^2$ | Intercept | Mean | CV |
| **Blueberry C:N top models** | | | | | | |
| 4 | 0.000 | −90.545 | 0.550 | -866.378 (±238.782) | 18.357 (±5.076) | 12.539 (±2.723) |
| 3 | 9.174 | −96.471 | 0.331 | -2.854 (±3.255) | | 12.121 (±3.255) |
| 3 | 14.716 | −99.242 | 0.196 | -806.460 (±312.860) | 17.364 (±6.654) | |
| 2 | 18.768 | −102.507 | 0.000 | 9.971 (±1.369) | | |
| **Blueberry N:P top models** | | | | | | |
| 2 | 0.000 | −102.507 | 0.000 | 9.971 (±1.369) | | |
| **Blueberry C:P top models** | | | | | | |
| 2 | 0.000 | −102.507 | 0.000 | 9.971 (±1.369) | | |
| **Red Maple N:P top models** | | | | | | |
| 2 | 0.000 | −102.507 | 0.000 | 9.971 (±1.369) | | |
| **Red Maple C:N top models** | | | | | | |
| 2 | 0.000 | −102.507 | 0.000 | 9.971 (±1.369) | | |

**Table S3:** Quantification of home range core area overlap for the four individuals with more than one year of telemetry sampling (see Section S5.1), as calculated with the Bhattacharyya affinity index (Fieberg and Kochanny, 2006; Winner et al., 2018). Note that we calculated the overlap for the whole Utilization Distribution (UD) of each individual between consecutive years of telemetry sampling. Figures S3 to S6 provide a visual representation of the overlap among these yearly home ranges.

| UD1 | UD2 | Bhattacharyya Index |
|---|---|---|
| **A1425** | | |
| 2018 | 2018 | 1.000 |
| 2019 | 2018 | 0.828 |
| 2018 | 2019 | 0.828 |
| 2019 | 2019 | 1.000 |
| **A1698** | | |
| 2018 | 2018 | 1.000 |
| 2019 | 2018 | 0.325 |
| 2018 | 2019 | 0.325 |
| 2019 | 2019 | 1.000 |
| **A3719** | | |
| 2017 | 2017 | 1.000 |
| 2018 | 2017 | 0.897 |
| 2017 | 2018 | 0.897 |
| 2018 | 2018 | 1.000 |
| **A3769** | | |
| 2018 | 2018 | 1.000 |
| 2019 | 2018 | 0.829 |
| 2018 | 2019 | 0.829 |
| 2019 | 2019 | 1.000 |

### S7 Supplementary Figures

Here we provide supporting visual materials for our field protocols, analyses, and results. Figure S1 show the location of our study area. For an interactive study area map, please see the Supporting Code document. Figure S2 provides a visualization of the grid layout. Figures S3 to S6 show the overlap in the 50%, 75%, and 90% aKDE isopleths of the four individuals for which we had multiple sampling years. Figure S7 shows a comparison of the C, N, P content of red maple and lowland blueberry in our hare study area.

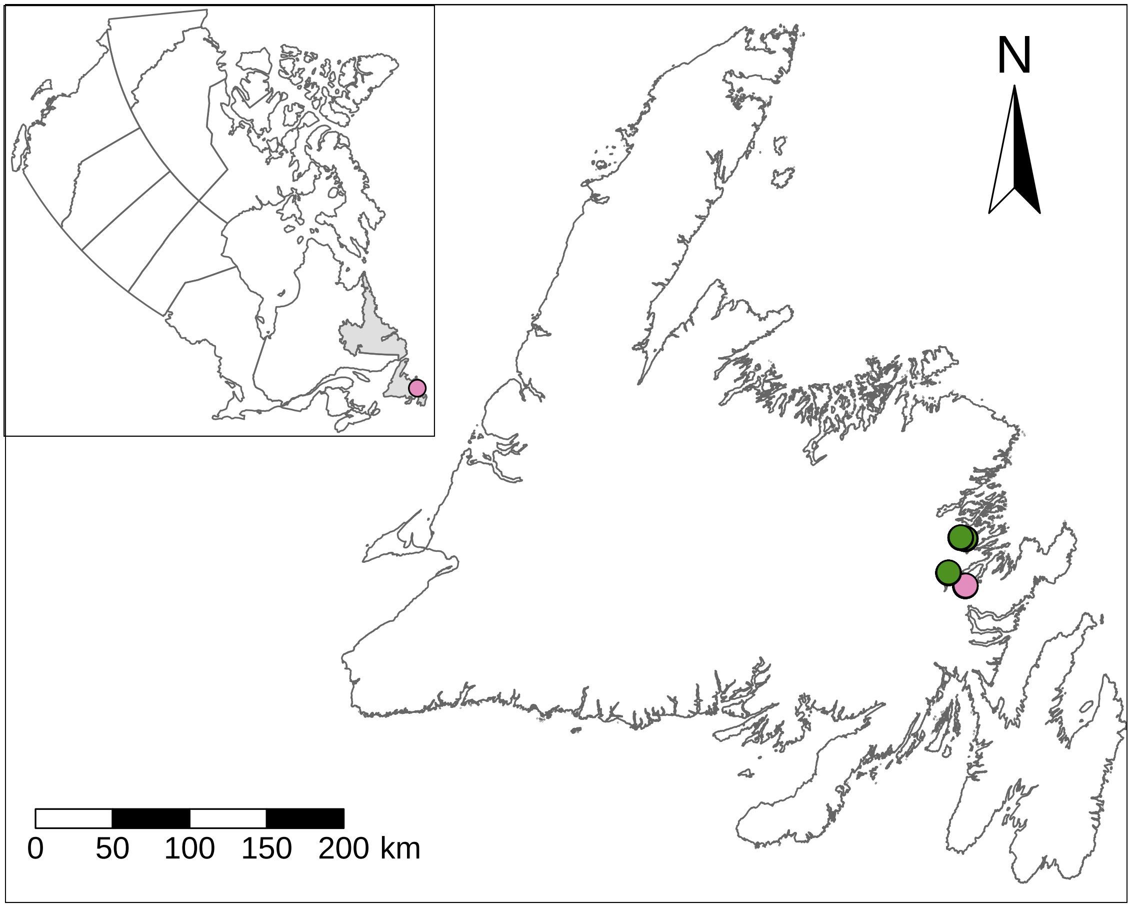

**Figure S1:** Map of study area. Each dot represents a sampling area. The pink dot represent the hare study area, where both plant sampling and snowshoe hare triangulation took place. The green dots show the location of the three areas where we only performed plant sampling. The inset shows the relative position of the island of Newfoundland to the rest of Canada, with the pink dot showing the location of our study area on the island of Newfoundland.

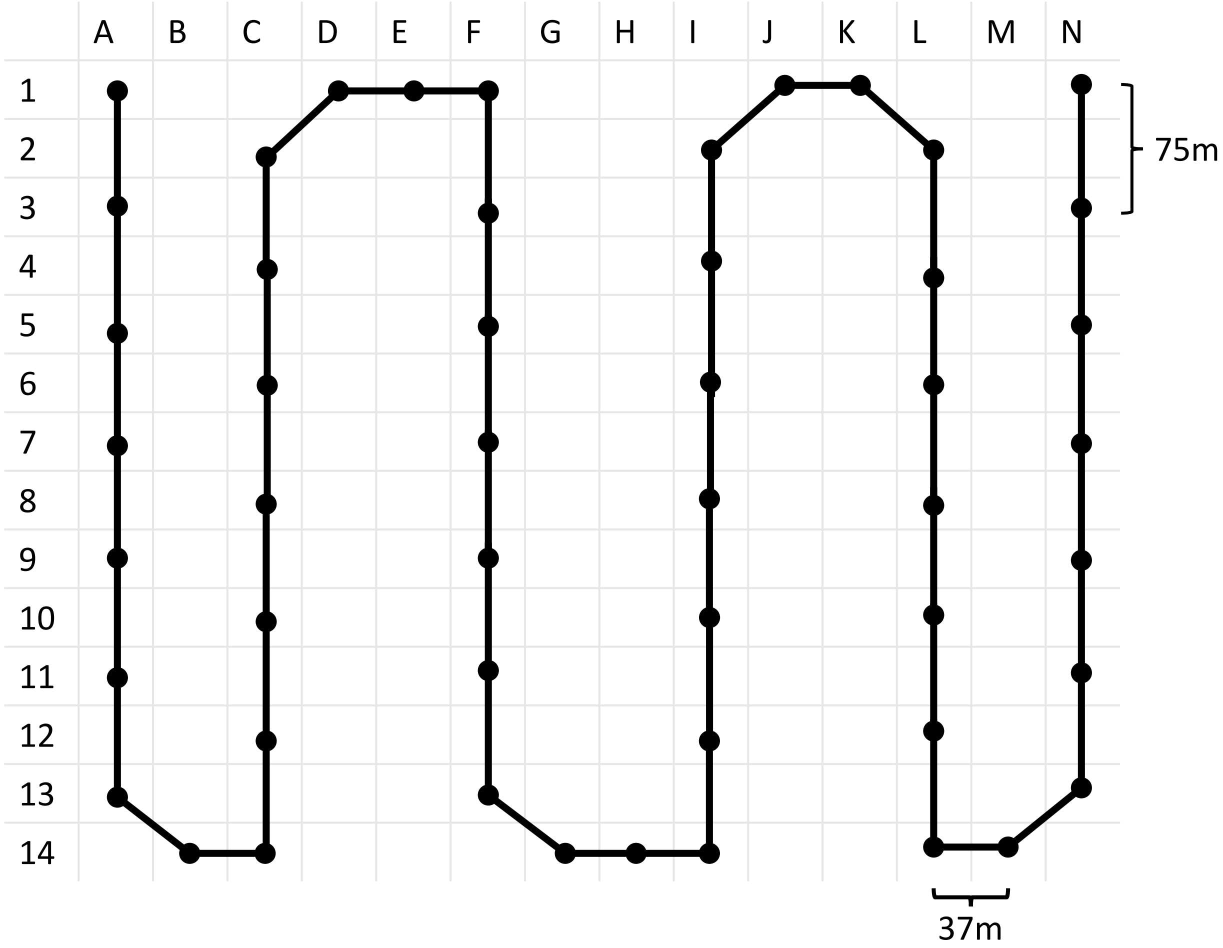

**Figure S2:** Layout of a snowshoe hare trapping grid. The grid consists of 14 rows and 14 columns (fine gray lines). At each intersection we placed a trap (full circles), alternating odd and even rows on adjacent lines. Traps on a line were 75 m apart, the only exceptions being traps placed on turns for which the distance was necessarily shorter.

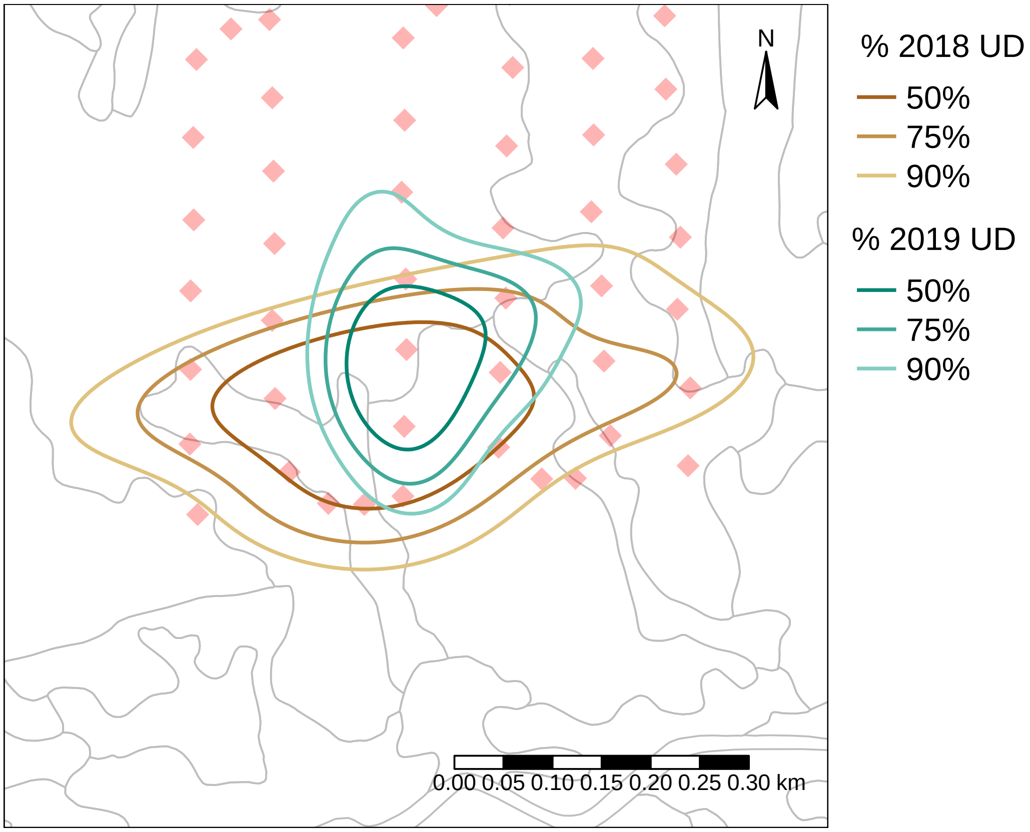

**Figure S3:** Map showing the high degree of overlap between 2018 and 2019 kernel Utilization Distribution estimates for snowshoe hare A1425. For both years, the map shows the relocations ($n = 26$ and $n = 33$, respectively), as well as the 50%, 75%, and 90% isopleth contours. Light red diamonds show traps locations on the trapping grid. Grey lines enclose Forest Resource Inventory-classified polygons.

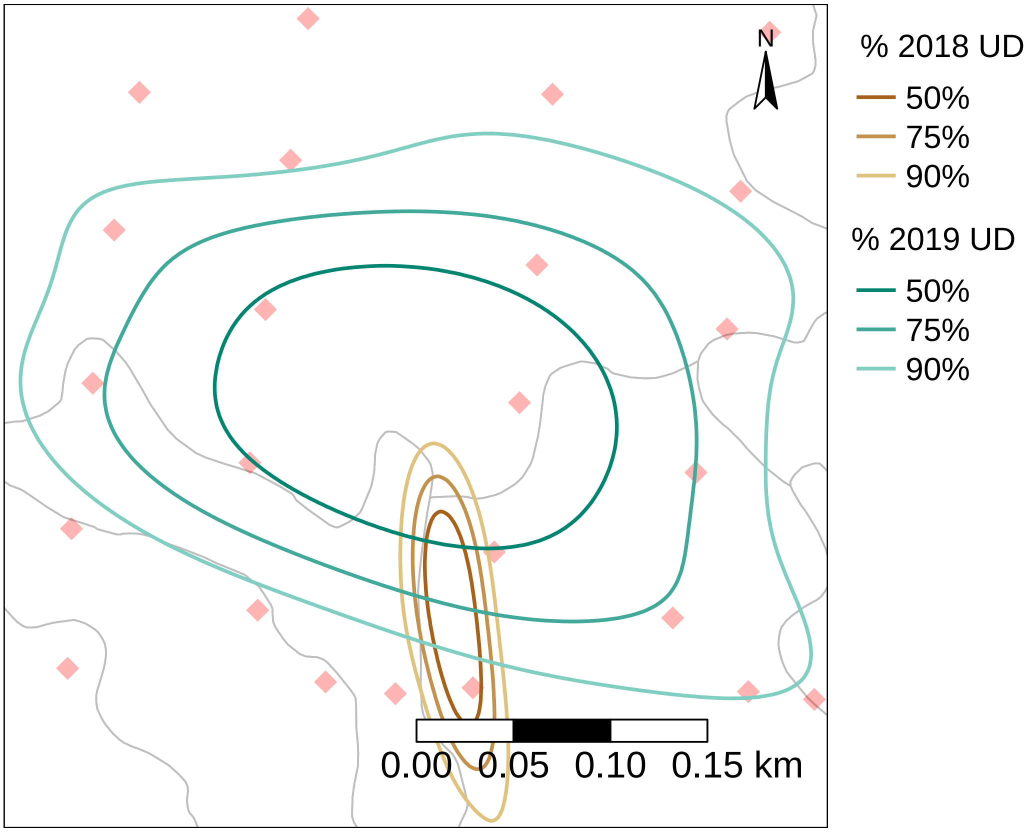

**Figure S4:** Map showing the overlap between 2018 and 2019 kernel Utilization Distribution estimates for snowshoe hare A1698. For both years, the map shows the relocations ($n = 28$ and $n = 31$, respectively), as well as the 50%, 75%, and 90% isopleth contours. All specifications as in Figure S3.

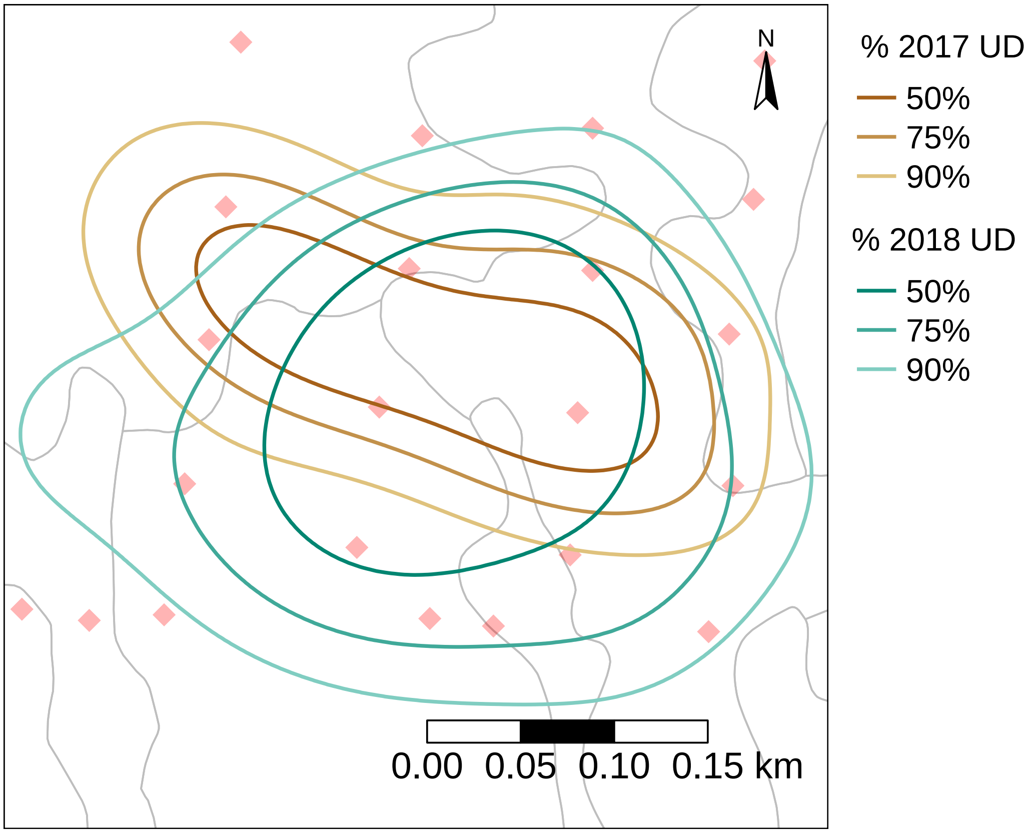

**Figure S5:** Map showing the high degree of overlap between 2017 and 2018 kernel Utilization Distribution estimates for snowshoe hare A3719. For both years, the map shows the relocations ($n$ = 36 and $n$ = 29, respectively), as well as the 50%, 75%, and 90% isopleth contours. Note that we do not show relocations or the aKDE for 2019 as this individual died halfway through the sampling season, when not enough relocations had been collected to produce an estimate of its UD. All specifications as in Figure S3.

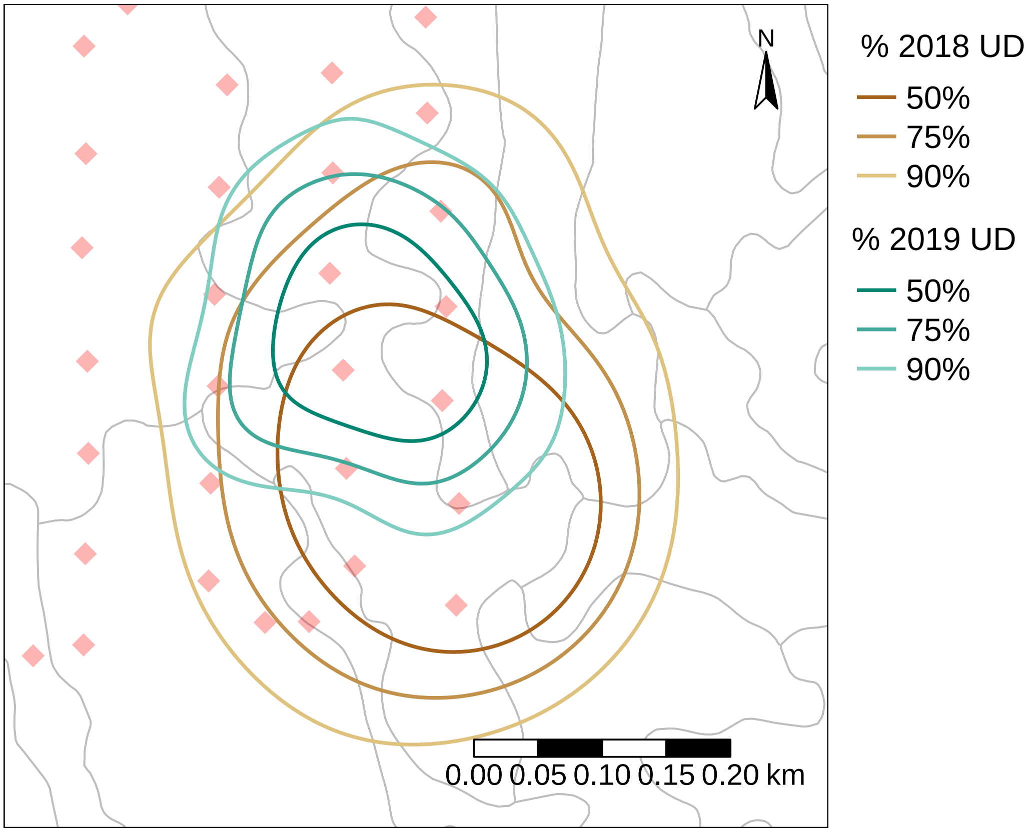

**Figure S6:** Map showing the high degree of overlap between 2018 and 2019 kernel Utilization Distribution estimates for snowshoe hare A3769. For both years, the map shows the relocations ($n$ = 26 and $n$ = 45, respectively), as well as the 50%, 75%, and 90% isopleth contours. All specifications as in Figure S3.

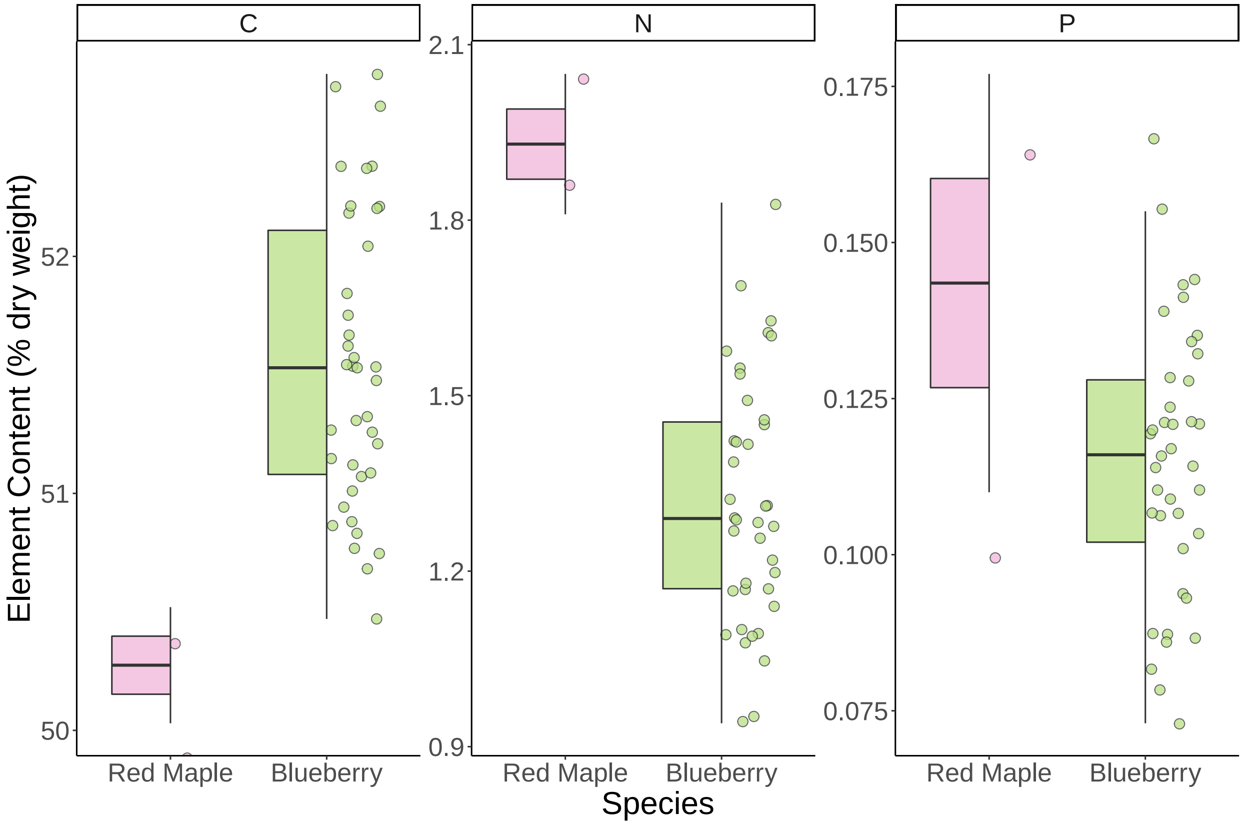

**Figure S7:** Comparison of the elemental content of C, N, and P in red maple and lowland blueberry in our hare study area. While red maple appears to have higher amounts of both N and P, the rarity of this plant on the grid makes lowland blueberry a much more profitable forage species for snowshoe hares.
